# Transcriptional Response to Tumor Mutational Burden Is Consistent Across Cancer Types

**DOI:** 10.64898/2026.08.27.744035

**Authors:** Karen Y. Shih, Onn Brandman, Monte M. Winslow, Dmitri A. Petrov

## Abstract

Tumor mutational burden (TMB) shapes tumor transcriptional state, but studies typically describe this response as an average effect pooled across cancer types. Whether that average reflects a consistent response present within individual cancer types, or is an artifact of merging heterogeneous, tissue-specific responses, remains unresolved. Here we analyze ∼9,100 tumors across 32 TCGA cancer types to test whether the transcriptional response to TMB is genuinely consistent across tissues. We construct a TMB axis score from TMB-associated genes upregulated with increasing TMB, yielding a sample-level measure of response strength, and subsequently decompose it at the component and pathway/complex levels. The pooled transcriptional response to TMB stays largely consistent within each cancer type, and no single cancer is driving the pooled signal. This consistency was also observed at the component and pathway/complex levels. These findings support TMB as a promising tissue-agnostic signature, with implications for tissue-agnostic therapeutic targeting.

**Summary:** Increasing tumor mutational burden (TMB) reflects the accumulation of mutations that impose a broad physiological burden on cancer cells, including increased DNA damage responses, protein misfolding, metabolic stress, and immune signaling. Here, we quantify TMB-associated transcriptional responses across and within cancer types to test whether this response is consistent. We construct a TMB axis score capturing genes upregulated with increasing TMB, generating a sample-level measure of response strength. Most cancer types exhibit a consistent response magnitude as TMB increases. TMB variance, more than sample size, constrains the detectability of this response within individual cancer types. Because baseline expression of TMB-associated genes differs across cancer types, pan-cancer comparisons can mask this signal. In contrast, within-cancer-type normalization reveals a transcriptional response spanning multiple cellular pathways that is consistent across cancer types despite differences in baseline gene expression.

## Introduction

Tumor mutational burden (TMB), defined as the number of somatic mutations per megabase of genome sequenced, spans several orders of magnitude across and within cancer types^1^. Mutations can arise through many different mechanisms, such as DNA replication errors, mutagen exposure, and deficient repair, which can vary across cancer types. Yet only a small fraction of the resulting mutations, driver mutations, are recurrent and have clear downstream cellular consequences^2^. The vast majority, passenger mutations, accumulate stochastically and are non-recurring among patients, regardless of cancer type^3^. TMB is overwhelmingly made up of passenger mutations^2^. The non-specificity of passengers predicts that their downstream cellular consequences might also be generic instead of tissue-specific^4^.

A previous pan-cancer analysis found that, on average, high TMB is associated with upregulation of multiple pathways, including proteostasis (e.g., the proteasome), DNA damage repair (e.g., homologous recombination), and transcriptional, metabolic, and immune signaling pathways (e.g., the spliceosome, carbon metabolism, and antigen processing and presentation)^4^. These transcriptional responses suggest that cells experience multifaceted cellular stress as TMB increases. However, it remains unclear whether the patterns of cellular transcriptional response to TMB hold across cancer types, or whether they vary strongly depending on the cancer-type baseline gene expression. Under the divergent response model, the transcriptional response to TMB might be heterogeneous across cancer types because baseline gene expression constrains, or obviates the need for, the response. Tumors that already express TMB-associated genes at high levels may experience diminishing returns from further upregulation, while others may be unresponsive to TMB—maintaining low TMB-associated gene expression regardless of the mutational burden, suggesting that either stress does not scale with TMB in these contexts or alternative buffering mechanisms are involved. Under this model, the observed pan-cancer association reflects the average of a diverse set of tissue-specific responses that differ in both direction and magnitude.

Under the consistent response model, baseline gene expression does not shape the response. Regardless of baseline gene expression, all tumor types upregulate nearly the same set of genes proportionally with increasing TMB. Distinguishing these models requires more than demonstrating a pan-cancer association because such an association is expected from any analysis that aggregates data across cancer types. The key test is whether the response is consistent within individual cancer types, such that the pan-cancer average reflects broadly shared biology rather than an aggregate over heterogeneous tissue-specific responses.

Here, we first construct a quantitative TMB axis, a low-dimensional representation of the transcriptional response associated with increasing TMB, rather than examining gene-level associations individually. Projecting the expression profile of each tumor onto this axis yields a sample-level measure of response strength that can be compared across and within cancer types. Using this framework, we test whether the TMB-associated transcriptional response is consistent across individual cancer types, and whether this consistency holds at increasingly granular levels. We further decompose the TMB axis into its proteostasis and non-proteostasis components, and each component into its constituent complexes and pathways. We additionally ask if apparent variability in the response across types reflects true biological differences or sampling artifacts— including differences in cohort size and TMB variance. Finally, we examine whether response magnitude depends on baseline expression levels of TMB-associated genes or is consistent across tissues independent of their expression baselines.

After accounting for between-cancer-type differences in baseline expression, our data show that the transcriptional response to increasing TMB, spanning multiple pathways, is broadly consistent across biologically distinct cancer types, indicating a shared cellular program downstream of tissue-specific biology. This consistency has direct implications for the use of TMB as a tissue-agnostic biomarker. TMB is already a predictive biomarker for immunotherapy response in solid tumors, supporting the tissue-agnostic FDA approval of pembrolizumab for TMB-high tumors^5^, an association typically attributed to increased neoantigen load^6^. Our finding of a broadly consistent transcriptional response suggests that the tissue-agnostic utility of TMB may extend beyond neoantigen-driven immunotherapy to additional biomarkers or therapeutic vulnerabilities.

## Results

### A transcriptional axis quantifies the response to tumor mutational burden

We defined tumor mutational burden (TMB) as the log_10_-transformed total number of point mutations in protein-coding genes, including nonsense, missense, and silent mutations. This definition follows our previous study^4^, which showed that different classes of point mutations are highly collinear and largely reflect a shared underlying process of mutation accumulation, with passenger mutations comprising the majority of somatic alterations^2,7^.

To quantify the transcriptional response associated with TMB at the sample level, we defined a TMB-associated transcriptional axis (hereafter referred to as the TMB axis) (Figure 1A). Using the TCGA Pan-Cancer gene expression RNA-seq data, we first identified genes whose expression was significantly associated with TMB using a gene-wise generalized linear mixed model (GLMM)^4,8^, which accounts for tumor purity and cancer-type differences. We focused on genes positively associated with TMB (β > 0, FDR < 0.05), since we aimed to capture transcriptional upregulation with elevated TMB.

**Figure 1.**
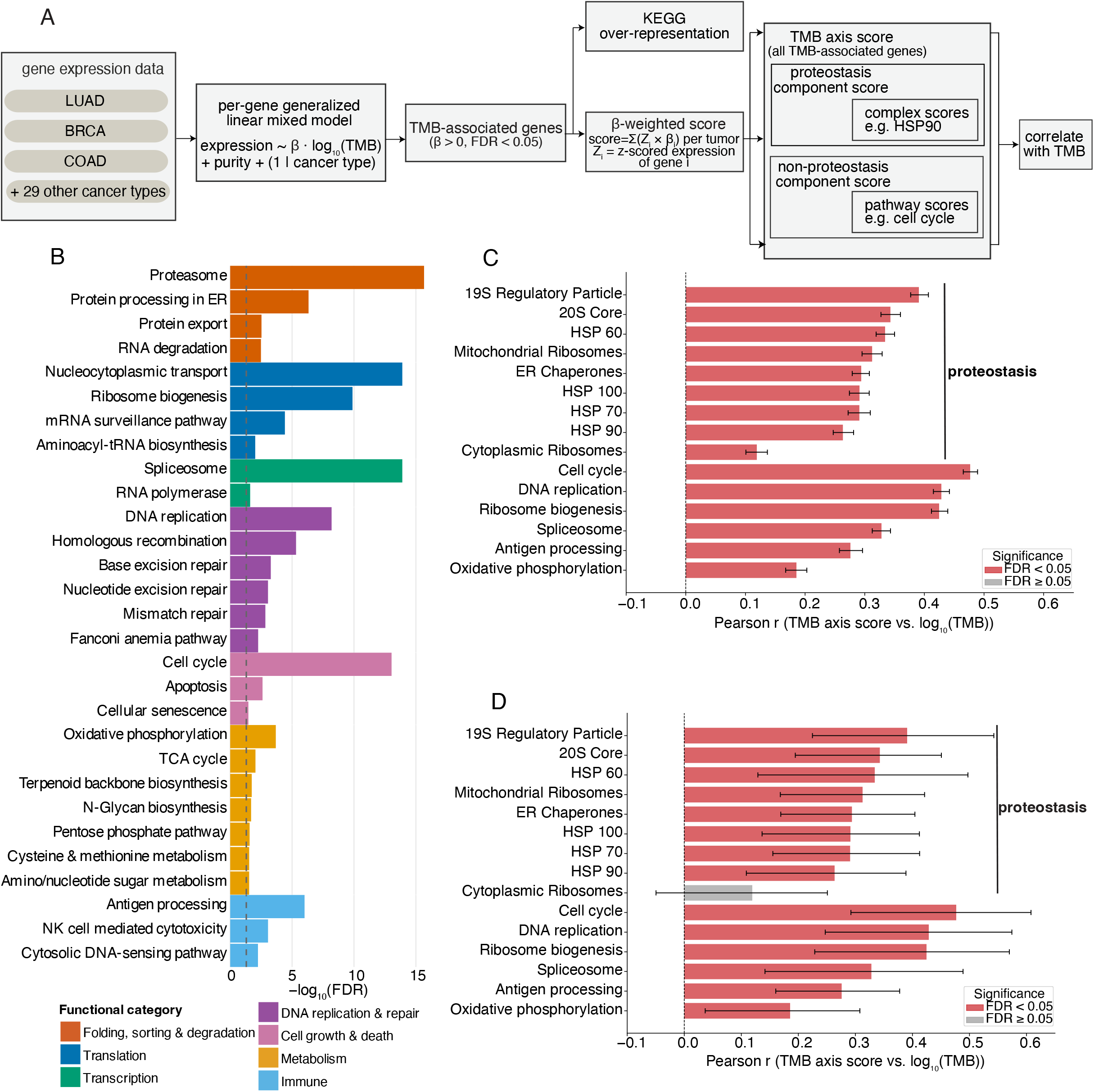
Overview of modeling and scoring framework. (A) Schematic overview of the modeling and tumor scoring framework. Gene expression data from 32 TCGA cancer types (e.g., LUAD, BRCA, COAD) were modeled using a generalized linear mixed model (GLMM): expression ∼ β · log_10_(TMB) + purity + (1 | cancer type). Per-gene regression coefficients (β) representing the association with TMB were extracted. All analyses were restricted to genes with positive coefficients (β > 0) and FDR < 0.05, corresponding to genes positively associated with TMB. Gene expression values were z-scored globally. KEGG was used for over-representation analysis. All scores were computed per sample as weighted sums of z-scored gene expression values (Z), using β as weights. Scores are computed at three nested levels: the TMB axis score, over all TMB-associated genes; component scores (proteostasis and non-proteostasis), which partition the axis score; and pathway or complex scores, over the individual functional gene sets within each component. (B) KEGG pathway over-representation analysis of all TMB-associated genes (β > 0, FDR < 0.05). Pathways are grouped into functional categories (e.g., folding, sorting & degradation, translation, transcription, DNA replication & repair, cell growth & death, metabolism, immune). The background gene set for enrichment analysis consisted of all genes tested in the GLMM. Pathways are colored by functional category (KEGG subcategory). Bar length represents enrichment significance (−log_10_(FDR)). Dashed line indicates FDR = 0.05. (C) Within-cancer-type bootstrap estimates of pathway/complex score–TMB correlations. Nine out of ten proteostasis complexes in the annotation and six representative non-proteostasis KEGG pathways are shown. Small heat shock proteins are omitted because no member passed the TMB-association threshold (β > 0, FDR < 0.05). Complex and pathway scores were computed per tumor sample and correlated with log_10_(TMB). Error bars are 95% bootstrap confidence intervals from 1,000 replicates in which tumors were resampled with replacement within each cancer type and the pan-cancer correlation recalculated. (D) Cancer-type bootstrap estimates of pathway/complex score–TMB correlations. The same complexes and pathways in (C) are shown here. Cancer types were resampled with replacement to assess the stability of pathway/complex score–TMB correlations across different cancer-type compositions. Complex and pathway scores were computed per tumor sample. Error bars represent 95% bootstrap confidence intervals from correlations recalculated across all tumors in the resampled dataset for each of the 1,000 cancer-type bootstrap replicates.

We then projected each tumor onto a one-dimensional transcriptional axis by computing a weighted sum of normalized gene expression values over TMB-associated genes using the estimated regression coefficients (β) as weights. Normalization places genes on a comparable scale, while β-weighting emphasizes genes most strongly associated with TMB. We refer to this metric as the TMB axis score: a continuous, per-sample summary of TMB-associated gene upregulation that reduces a high-dimensional transcriptional response to a single value.

Using the same projection framework, we partitioned the TMB axis score into proteostasis and non-proteostasis components by restricting the weighted sum to proteostasis genes and to all remaining TMB-associated genes, respectively. Within the proteostasis component, we scored curated protein complexes^9,10^ (e.g., the HSP90 chaperone complex): for each complex, the score used its member genes that were significantly TMB-associated (β > 0, FDR < 0.05), weighted by their regression coefficients, yielding a complex score. For the non-proteostasis component, we instead grouped the significant TMB-associated genes by KEGG pathway (e.g., cell cycle) and scored each pathway from its member genes, yielding a pathway score.

Functional enrichment analysis over all TMB-associated genes identified enriched categories encompassing protein folding, sorting and degradation^4,11^, alongside cell growth and death, DNA replication and repair, metabolism, and immune signaling (Figure 1B). These results suggest that increasing TMB is associated with a transcriptional response spanning diverse cellular pathways.

Across chaperone and proteasome complexes, the fraction of genes significantly upregulated with TMB varied markedly, from none (small heat shock proteins, small HS) to all members (19S regulatory particle, 20S core), indicating the response is not uniform across complexes (Figure 1—figure supplement 1A).

Next, we decomposed the TMB axis into proteostasis and non-proteostasis components and scored their constituent complexes and pathways. Both sets exhibited broadly positive associations with TMB, stable under within-cancer-type bootstrapping (Figure 1C; Figure 1— figure supplement 1B), as expected given that the TMB axis is defined by positively associated genes. Correlation magnitudes varied across functional categories (r = 0.12–0.54).

To assess the robustness of the pan-cancer observation in Figure 1C to cancer-type composition, we performed cancer-type bootstrapping to estimate between-cancer variability, repeatedly resampling with replacement the 32 TCGA cancer types and recalculating the pan-cancer correlation between the sample-level pathway scores and log_10_(TMB) values (Figure 1D). Except for cytoplasmic ribosomes, all other proteostasis complexes and non-proteostasis pathways remained positively correlated with increasing TMB (Figure 1D; Figure 1—figure supplement 1C). In a complementary leave-one-cancer-type-out (LOCO) analysis, we held the TMB-associated gene set fixed and re-estimated β on all but one cancer type, then projected the held-out type onto the TMB axis and correlated its TMB axis scores with log_10_(TMB). The LOCO validation correlations closely matched those obtained using the full dataset, indicating that no single cancer type drives the axis structure and that the underlying transcriptional signal is broadly shared across tumor types (Figure 1—figure supplement 1D). Although a small number of cancer types showed weak or negative TMB axis score–TMB correlations in both the full and LOCO models, the two estimates were in near-perfect agreement across the full range (Pearson r = 0.996), demonstrating that β values are not meaningfully influenced by the inclusion or exclusion of any single cancer type.

To examine whether the TMB-axis decomposition generalizes beyond the samples used to estimate gene regression coefficients, we held out 30% of tumors within each cancer type (gene set fixed on the full dataset; only β refit), fit the model to the remaining 70%, and computed pan-cancer Pearson correlations between axis scores and log_10_(TMB) on the validation set. The validation correlations closely matched those obtained using the full dataset (full dataset vs. validation—TMB axis: r = 0.55 vs. 0.54; proteostasis component: r = 0.39 vs. 0.37; non-proteostasis component: r = 0.55 vs. 0.54), confirming that the axis scores are not an artifact of overfitting to the full sample. Across both the full-dataset and validation analyses, the correlation was driven primarily by the non-proteostasis component, with the weaker proteostasis association possibly reflecting its smaller gene set.

### A consistent transcriptional response to TMB across cancer types

To characterize the per-cancer transcriptional response associated with TMB, we computed per-sample TMB axis scores and assessed their relationship with TMB within individual cancer types. Per-cancer correlations, ordered by the signed values of the TMB axis score–TMB correlation, were, by design, predominantly positive across cancer types for the overall TMB axis as well as its proteostasis and non-proteostasis components (Figure 2A; Figure 2—figure supplement 1A, B). 29 of 32 cancer types showed a positive point estimate for the correlation between TMB axis score and TMB (sign test, p = 2.6 × 10^− 6^; Figure 2A), with KIRP, UVM, and THYM being the only cancer types with a negative point estimate. Based on the divergent response model, we would expect heterogeneous correlations with non-overlapping confidence intervals across cancer types, because the transcriptional response would be cancer-type-specific. Instead, we observed substantial overlap of within-cancer-type bootstrap confidence intervals, indicating that the magnitude of the TMB-associated transcriptional response is consistent across most cancer types. This similarity in magnitude supports the consistent response model, in which the pan-cancer average reflects broadly shared biology rather than an aggregate over heterogeneous tissue-specific responses.

**Figure 2.**
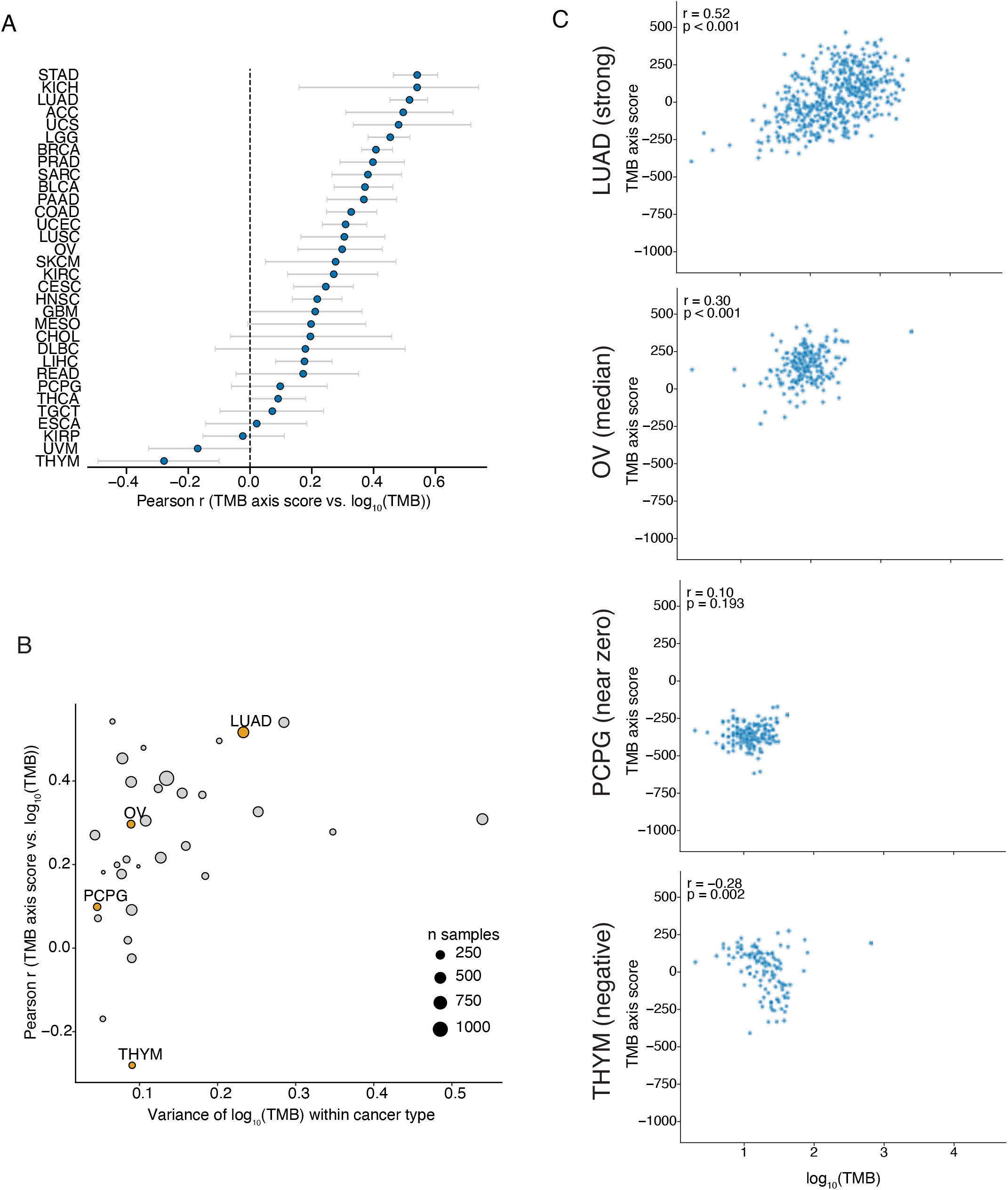
Consistent transcriptional response to TMB across cancer types. (A) Summary of correlations between log_10_(TMB) and TMB axis scores across cancer types. Each point represents the Pearson correlation within a single cancer type. Error bars represent 95% bootstrap confidence intervals obtained by repeatedly resampling tumors within each cancer type and recalculating the correlation. Most cancer types exhibit positive associations, and the overlapping confidence intervals support a consistent transcriptional response to increasing TMB across cancer types. (B) Influence of data structure on the detectability of the TMB-associated transcriptional response. Each point represents a cancer type, with the x-axis showing variance of log_10_(TMB) and the y-axis showing the corresponding Pearson correlation with the TMB axis. Point size reflects sample size. Cancer types with lower TMB variance tend to show weaker correlations, indicating that apparent differences in correlation are driven in part by TMB variance. Highlighted cancer types (LUAD, OV, PCPG, THYM) are examined individually in (C). (C) Relationship between log_10_(TMB) and TMB axis score across representative cancer types. Each point represents a tumor. Representative cancer types were selected to illustrate a range of observed correlations, including strong positive (top ∼10%), typical (median), near-zero, and negative. Cancer types with near-zero or negative correlations exhibit a relatively narrow range of TMB values, illustrating how limited TMB variance can reduce the apparent strength of the association. LUAD, lung adenocarcinoma; OV, ovarian serous cystadenocarcinoma; PCPG, pheochromocytoma and paraganglioma; THYM, thymoma.

Yet, the observed per-cancer correlation strength varied, in part due to differences in dataset structure. To quantify the contribution of dataset structure to variability in between-cancer-type correlations, we tested associations between per-cancer TMB axis score–TMB correlations and two dataset-level factors—TMB variance and sample size (Figure 2B). Across 32 cancer types, TMB variance was the stronger predictor (Spearman ρ = 0.45, p = 0.011), whereas sample size was not significantly associated (Spearman ρ = 0.28, p = 0.12). Low TMB variance within a cancer type may reflect either limited variation in the sampled cohort or an intrinsic property of that tumor type, in which mutational burdens naturally fall within a relatively narrow range.

To illustrate the range of observed associations, we examined four representative cancer types spanning strong positive, typical, near-zero, and negative correlations. Lung adenocarcinoma (LUAD), which exhibits one of the broadest TMB distributions (variance = 0.23; fourth highest of 32 cancer types), showed one of the strongest positive correlations. Ovarian serous cystadenocarcinoma (OV; variance = 0.09; 11th lowest of 32) displayed a correlation near the pan-cancer median. Pheochromocytoma and paraganglioma (PCPG; variance = 0.05; second lowest of 32) showed little association, consistent with its restricted TMB variance. In contrast, thymoma (THYM; variance = 0.09) exhibited a negative correlation despite comparable TMB variance to OV (Figure 2C; Figure 2—figure supplement 1C), suggesting that factors beyond TMB variance may also contribute in some cancer types.

We further quantified the magnitude of the transcriptional response within each cancer type by estimating slopes while controlling for tumor purity. Slopes were consistent across cancer types for the overall TMB axis and for proteostasis and non-proteostasis components, with broadly overlapping confidence intervals (Table S1). This consistency extended to individual proteostasis complexes and non-proteostasis pathways (Table S2), indicating that the transcriptional response to TMB is a shared feature across distinct biological programs.

### Cancer-type-specific baseline expression masks consistent TMB responses

To compare the TMB-associated transcriptional response within individual cancer types, we separated tumors into low- and high-TMB groups using the median TMB of each cancer type. Although high-TMB tumors consistently exhibited higher axis scores than low-TMB tumors within each cancer type, the absolute score range varied substantially across cancer types (Figure 3A). For example, stomach adenocarcinoma (STAD) tumors exhibited substantially higher baseline axis scores than adrenocortical carcinoma (ACC) tumors (Figure 3—figure supplement 1A), reflecting higher average expression of TMB-associated genes in STAD independent of mutational burden. Such baseline differences across cancer types could confound direct comparisons of TMB-associated responses.

**Figure 3.**
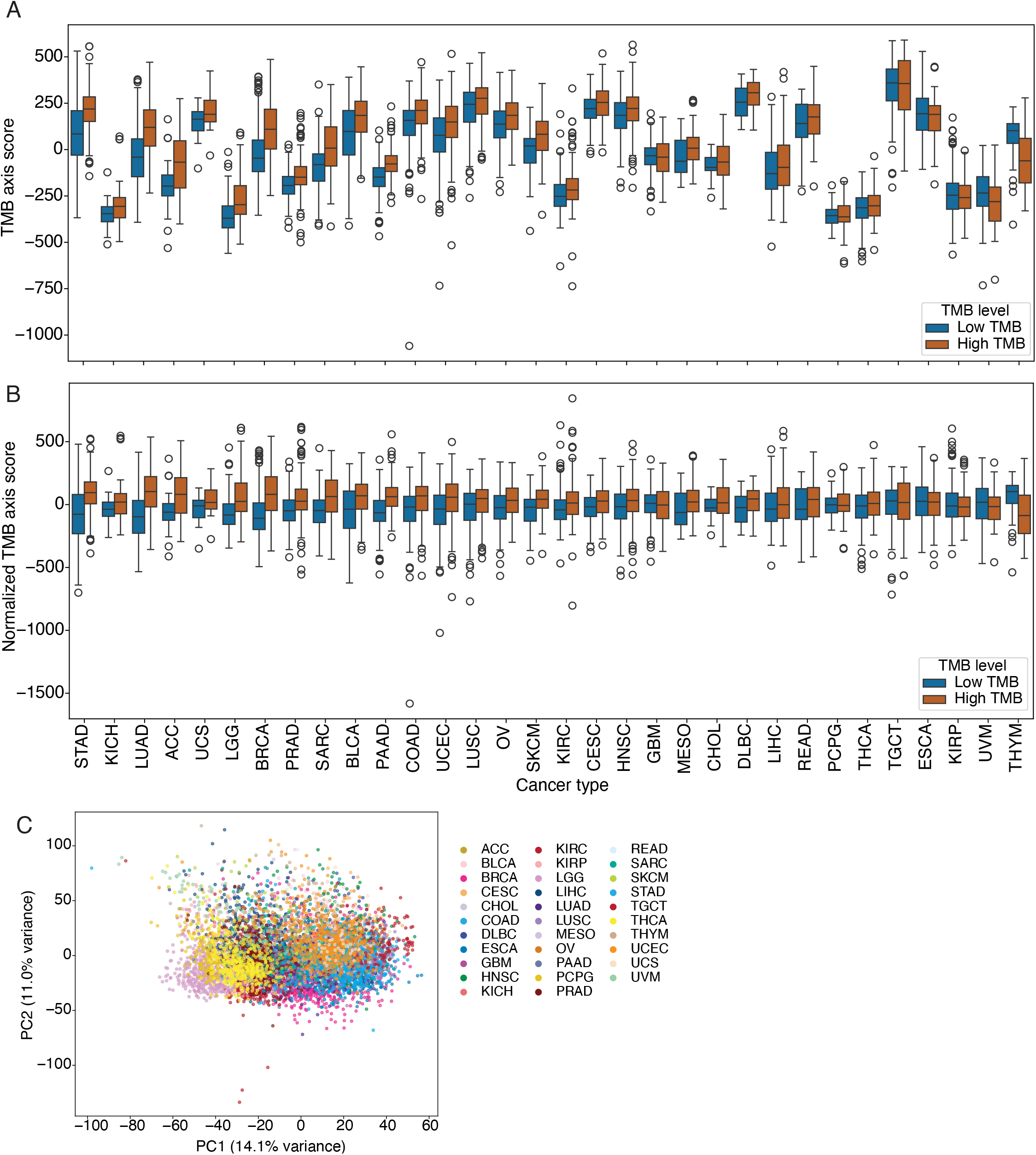
Within-cancer normalization reveals a consistent TMB-associated transcriptional response. (A) TMB axis scores computed from globally gene-wise z-scored expression across TCGA samples, stratified by low- and high-TMB groups within each cancer type. Cancer types exhibit widely different baseline axis scores, reflecting global transcriptional differences between tumor types independent of mutational burden. Although high-TMB tumors tend to score higher than low-TMB tumors in most cancer types, the large variation in absolute scores across cancer types obscures direct comparison of TMB responses. (B) TMB axis scores after within-cancer gene-wise z-scored normalization. Normalizing within each cancer type removes baseline offsets, placing all cancer types on a comparable scale. Under this normalization, high-TMB tumors consistently exhibit higher axis scores than low-TMB tumors in most cancer types, revealing a broadly consistent transcriptional response to increasing mutational burden. (C) PCA of TMB-associated gene expression across TCGA samples, colored by cancer type. Despite restricting the analysis to TMB-associated genes, samples cluster primarily by cancer type, indicating that cancer-type-specific transcriptional structure dominates global expression variation and underscoring the necessity of within-cancer-type normalization for detecting TMB-associated responses.

To account for cancer-type-specific baseline expression, we standardized gene expression within each cancer type prior to projection onto the TMB axis. Following within-cancer-type normalization, high-TMB tumors consistently exhibited higher TMB axis scores than low-TMB tumors in all but three cancer types (KIRP, UVM, and THYM—the same three with negative TMB axis score–TMB correlations; Figure 3B), highlighting a consistent transcriptional response to increasing TMB.

The importance of within-cancer-type normalization is further supported by principal component analysis of TMB-associated genes, which revealed that tumors clustered primarily by cancer type despite restricting the analysis to TMB-associated genes (Figure 3C). Similar clustering was observed for proteostasis and non-proteostasis gene subsets (Figure 3—figure supplement 1B), indicating that cancer-type-specific transcriptional structure remains a dominant source of variation even within TMB-associated programs.

In summary, cancer-type-specific expression variation can mask the transcriptional response to increasing TMB prior to within-cancer-type normalization. Once normalized, the response is broadly consistent—high-TMB tumors consistently score higher on the TMB axis than low-TMB tumors across cancer types. Notably, we observed neither the diminishing returns predicted for cancer types with high baseline expression nor the unresponsiveness predicted for those with low baseline expression, arguing against the divergent response model and in favor of a consistent transcriptional response to increasing mutational burden.

## Discussion

Here, we assessed consistency of the TMB-associated transcriptional response between and within cancer types. We aimed to distinguish two response models: a consistent response model, in which cancer types share the same proportional response to increasing TMB despite differing baselines, and a divergent response model, in which the response itself differs across cancer types. Because our GLMM is designed to detect a shared effect across cancer types, it estimates a single slope representing the average within-cancer-type relationship between TMB and expression for each gene. The TMB axis is then defined from genes with a positive average slope, meaning that a positive pan-cancer association is guaranteed by construction and is therefore uninformative about cross-cancer-type consistency on its own. The key question this leaves open is whether the axis captures a genuinely shared response across cancer types or merely averages over heterogeneous, type-specific effects. These possibilities can be distinguished: a cancer type with no response would exhibit a near-zero within-cancer-type correlation, whereas a cancer type with a reversed response would exhibit a negative correlation. Neither pattern is apparent from the fixed-effect estimates alone.

Our data support the consistent response model. Within-cancer-type correlations between TMB and axis scores were mostly consistent in magnitude across cancer types, with overlapping bootstrap confidence intervals. Robustness analyses (LOCO, cancer-type bootstrapping) confirmed the trend was not driven by any single type or subset. This consistency was not apparent until we accounted for the differences in baseline expression between cancer types because tissue of origin dominates absolute expression and masks the signal. Much of the variability in correlation strength reflected differences in TMB variance rather than differences in the underlying transcriptional response. We did not observe the diminishing returns and unresponsiveness predicted by the divergent response model. Baseline expression was unrelated to the response magnitude. Together, these results support a consistent TMB-associated transcriptional program across tissue contexts.

Why does this response remain consistent across such biologically diverse cancer types? Different cancer types might accumulate mutations through different mechanisms, but TMB summarizes their cumulative outcome. TMB is overwhelmingly composed of passenger mutations—roughly 97% of somatic mutations in tumors are passenger mutations, with only 2–6 driver mutations per tumor despite hundreds to thousands of total protein-coding mutations^2,12^. Passenger mutations are individually mild (each ∼100 times weaker than a driver) but collectively damaging, distributed across the proteome rather than concentrated in specific pathways^2,13^. The thermodynamic stability effects of mutations follow a similar distribution across proteins^14^. Rather than acting in isolation, the hundreds to thousands of passenger mutations collectively destabilize the proteome^4^. If the effect of specific mutated proteins outweighed the effect of the collective damage of the passenger mutations, we would have expected to see more cancer-type-specific responses to increasing TMB. Instead, all cell types experience generic stresses (proteotoxic, genotoxic, metabolic) rather than tissue-specific pathway activation. These stresses are sensed and responded to by conserved cellular programs, such as the proteostasis network, DNA damage response, and innate immune signaling^11,15^, that are constitutively present in all cell types.

There are several lines of direct evidence for the collective action of the passenger mutations. For instance, passenger mutations are linked through the Hill–Robertson interference process^16^. These mutations then accumulate during tumorigenesis and often scatter throughout the patients’ genomes and are non-recurring among patients^3^. Despite the deleterious passenger mutations, the tumors still arise through the tug-of-war with the rare but strongly advantageous drivers. This process is informative because unlike driver mutations, which are recurrent and gene-specific as they are under positive selection, passenger mutations do not exert their effect through the identity of the genes they hit but through their overwhelming number (∼100 times more numerous), in support of an aggregate process^13^. One phenotypic consequence of this cumulative passenger mutation burden is protein misfolding stress, shown by upregulation of proteostasis genes in high-TMB tumors; these tumors are correspondingly vulnerable to HSP90 and proteasome inhibitors^4^. Because this response depends on aggregate mutation count rather than which specific genes are hit, TMB, as a measure blind to mutation identity, is sufficient to capture it.

Our findings extend earlier work showing that high-TMB tumors upregulate proteostasis machinery^4^. That study focused on identifying TMB-responsive proteostasis complexes and establishing the functional importance of proteostasis for high-TMB cell viability, largely through pan-cancer analyses. This left open another question: whether the transcriptional response to TMB is consistent within individual cancer types, or whether pan-cancer signals aggregate over heterogeneous, tissue-specific responses. Because both a shared response and an average over divergent, cancer-type-specific responses can generate similar pan-cancer associations, distinguishing the two requires explicitly modeling within-cancer-type responses while controlling for tissue-of-origin baseline expression—the gap that motivated our approach. This distinction matters because only the former establishes that the response reflects a general feature of tumor biology rather than a statistical average, strengthening the rationale for tissue-agnostic therapeutic strategies.

The linear modeling used in our work may have missed other modes of TMB-associated regulation. Some cancers may experience non-linear responses such that certain pathways are activated only when TMB passes a certain threshold whereas others may saturate once TMB is sufficiently high. These additional modes could give us a more complete understanding of cellular responses to TMB-induced stresses and inform clinical interventions.

While we focused on the transcriptional response to increasing TMB in this study, proteomic and metabolomic data of tumors across the TMB spectrum could reveal downstream functional consequences of mutational accumulation not captured at the RNA level, potentially exposing additional therapeutic vulnerabilities. Existing genetic models of high mutation, including *POLE* proofreading-deficient^17,18^ and mismatch-repair-deficient^19,20^ mice and organoids, generate tumors with elevated TMB but have not been systematically characterized for the transcriptional response identified here. If these models recapitulate both the TMB-associated transcriptional response and the resulting dependence on proteostasis, they would directly support the proposed model linking TMB to proteostasis vulnerability and provide a platform for preclinical testing.

TMB is usually treated as a summary count of mutations. Our results suggest it can be viewed as a signature for the downstream cellular consequences of the mutation accumulation process, actively shaping the transcriptional changes in tumors. Although tissue of origin determines a tumor’s baseline gene expression, increasing TMB elicits a similar transcriptional response across cancer types. That shared response spans proteostasis, DNA repair, as well as metabolic and immune signaling pathways, suggesting that high-TMB tumors rely on a conserved set of stress response machinery. Our work reinforces downstream effects of TMB as a potential generic therapeutic vulnerability.

## Methods

### Data processing

TCGA pan-cancer RNA-seq (EB++ batch-corrected RSEM) and mutation calls were downloaded from the UCSC Xena PanCan Atlas (October 2025). Sample barcodes were truncated to 15 characters to harmonize expression, mutation, and purity tables. From 11,069 expression-matrix rows (11,060 unique barcodes), we retained primary solid tumors (sample_type_id == 1; n = 9,704), which excluded LAML (n = 173), as its samples are primary blood-derived cancers rather than solid tumors. Tumor mutational burden (TMB) was calculated from the TCGA MC3 public MAF (mc3.v0.2.8.PUBLIC.maf.gz) by counting missense, nonsense, and silent mutations per sample; because MC3 applies a uniform capture across samples, counts are directly comparable and were not normalized to mutations per megabase. Analyses used log_10_(mutation count + 1) as the continuous TMB covariate. An inner join with MAF-derived TMB excluded 600 primary-tumor rows lacking mutation counts, yielding a final cohort of 9,104 samples across 32 cancer types. ABSOLUTE tumor purity was joined to this cohort, and missing values were median-imputed within cancer type (314 values imputed; no samples were dropped for missing purity).

Before analysis, low-variance genes (variance ≤ 0.1 across samples) were removed, leaving 19,599 genes, and expression was z-scored per gene. TMB-associated axis genes were defined as genes positively associated (β > 0) with log_10_(mutation count + 1) in a per-gene linear mixed model with cancer type as a random intercept and tumor purity as a fixed effect (BH-adjusted FDR < 0.05; 4,903 genes). The same 4,903-gene symbol list was used for axis scoring, leave-one-cancer-out (LOCO) analyses, and train/validation analyses.

### Pan-cancer gene-level association with TMB

To identify genes associated with tumor mutational burden (TMB), we followed a framework described in our previous work^4^. TMB was calculated as the total number of missense, nonsense, and silent mutations per sample, consistent with the convention used previously^4^. All analyses using TMB as a continuous variable were performed using log_10_-transformed TMB values.

Briefly, we fit gene-wise GLMMs using the lme4 package in R. For each gene, expression values were normalized (z-scored) across all samples to place genes on a common scale and enable comparison of effect sizes. Models were specified as:

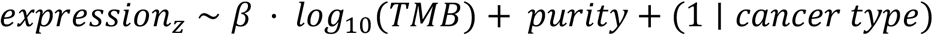

where β is the regression coefficient for log_10_(TMB) and cancer type was included as a random intercept. Downstream analyses used genes significantly and positively associated with TMB (β > 0, FDR < 0.05).

p-values and 95% confidence intervals for the association between gene expression and TMB (β for log_10_(TMB)) were obtained from the fitted models; p-values were adjusted for multiple hypothesis testing using the Benjamini–Hochberg procedure to control the false discovery rate (FDR).

### Gene set definitions

Proteostasis gene sets were defined at the protein level, comprising molecular chaperones, proteasomal subunits, and the ribosome, and totaled 265 unique gene symbols across 10 complexes. Chaperone complex genes were obtained from Hadizadeh Esfahani et al.^10^; co-chaperones were excluded. Proteasomal and ribosomal complex genes were obtained from CORUM^9^. For all proteostasis scores, only genes present in both the corresponding gene set and the TMB-associated set (β > 0, FDR < 0.05) were included; genes in a defined complex that were not significantly associated with TMB were excluded. This yielded 141 of the 265 proteostasis genes for scoring.

The non-proteostasis component comprised the remaining 4,762 TMB-associated genes (4,903 total minus 141 proteostasis genes). Separately, pathway gene sets corresponded to KEGG pathways defined directly from the TMB-associated genes; because these were derived from the TMB-associated genes, all pathway genes were TMB-associated by construction.

Note that the proteostasis gene set used for scoring (curated chaperones, proteasome subunits, and ribosome) is distinct from the KEGG “Folding, sorting and degradation” category used in the enrichment analysis; the two are defined from different sources and are not identical.

### Functional enrichment analysis

Over-representation analysis of the TMB-associated transcriptional program was performed using genes significantly upregulated with mutational burden in the pan-cancer GLMM (β > 0, FDR < 0.05). KEGG pathway annotations were used for functional enrichment analysis^21^.

Pathways annotated under the KEGG category “Human Diseases”, as well as pathways within the “Global and overview maps”, “Transport and catabolism”, and “Endocrine system” subcategories, were excluded from downstream analyses. The pathways “Oocyte meiosis” and “Progesterone-mediated oocyte maturation” were additionally excluded. These terms were excluded to emphasize broadly interpretable cellular processes and reduce redundancy from highly overlapping or disease-specific annotations.

For visualization and interpretation, several KEGG pathway and category names were shortened for display. Specifically, “Protein processing in endoplasmic reticulum” was relabeled as “Protein processing in ER”, “Antigen processing and presentation” was relabeled as “Antigen processing”, “Natural killer cell mediated cytotoxicity” was relabeled as “NK cell mediated cytotoxicity”, “Amino sugar and nucleotide sugar metabolism” was relabeled as “Amino/nucleotide sugar metabolism”, “Cysteine and methionine metabolism” was relabeled as “Cysteine & methionine metabolism”, “Citrate cycle (TCA cycle)” was relabeled as “TCA cycle”, “Folding, sorting and degradation” was relabeled as “Folding, sorting & degradation”, “DNA replication and repair” was relabeled as “DNA replication & repair”, and “Cell growth and death” was relabeled as “Cell growth & death”. All remaining pathways and categories retained their original KEGG pathway names.

### TMB axis score calculation

To summarize the transcriptional response to TMB at the sample level, we constructed a TMB-associated transcriptional axis (TMB axis). Using genes significantly positively associated with TMB (β > 0, FDR < 0.05), we computed a per-sample score (TMB axis score) as a weighted sum of normalized gene expression values, where weights correspond to the estimated GLMM regression coefficients β:

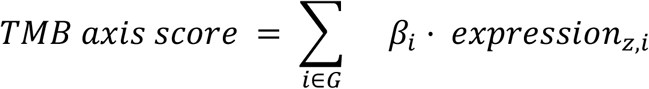

where expression_z,i_ is the normalized expression of gene i, and β_i_ is its estimated association with TMB, and *G* is the set of genes satisfying β_i_ > 0 and FDR < 0.05. Unless otherwise specified, gene expression values were gene-wise globally z-scored prior to score calculation. For analyses focused on within-cancer-type relative transcriptional changes, gene expression values were gene-wise z-scored within each cancer type before score calculation.

The TMB axis score decomposes into a nested hierarchy. It first partitions into proteostasis and non-proteostasis component scores, which together span G: the proteostasis component uses the subset of G in the proteostasis gene set (see Gene set definitions), and the non-proteostasis component uses the remaining genes in G. Within each component, finer-grained scores were computed for individual gene sets—complex scores within the proteostasis component and pathway scores within the non-proteostasis component—using the same weighted-sum formula restricted to the corresponding genes.

### Empirical null comparison

To assess whether observed gene-level associations exceeded those expected under a null model, we followed an empirical null framework described previously^4^. Specifically, we generated a permuted dataset in which log_10_-transformed TMB values were randomly shuffled within each cancer type, preserving cancer-type-specific distributions while disrupting associations with gene expression. The gene-wise GLMM was refit on the permuted data, and the same significance criteria (β > 0, BH FDR < 0.05) were applied. The fraction of significant genes within each proteostasis complex was then recomputed and compared with the observed fractions (Figure 1—figure supplement 1A).

### Quantification of TMB-associated transcriptional response

To quantify the relationship between TMB-associated transcriptional scores and mutational burden within cancer types, Pearson correlations between transcriptional scores and log_10_(TMB) were calculated globally across all tumors and separately for each cancer type.

95% bootstrapped confidence intervals and p-values were estimated from 1,000 bootstrap resamples unless otherwise specified. For within-cancer-type bootstrapping, tumors were resampled with replacement within each cancer type. For cancer-type bootstrapping, cancer types were resampled with replacement, and the statistic was recalculated for each bootstrap replicate. Two-sided p-values were derived from the bootstrap distribution as twice the smaller tail probability relative to zero correlation.

The magnitude of transcriptional response within each cancer type was estimated using linear regression models of the form:

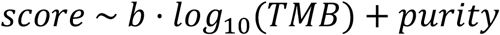

where the score is one of the TMB axis, proteostasis/non-proteostasis component scores, or pathway/complex scores, depending on the analysis. The slope b indicates the strength of the transcriptional response to increasing TMB. 95% confidence intervals were derived from the regression standard error.

Pathway scores in Figure 1C, D were computed for six KEGG pathways selected from the over-representation analysis: cell cycle (hsa04110, n = 97), DNA replication (hsa03030, n = 31), ribosome biogenesis (hsa03008, n = 51), spliceosome (hsa03040, n = 84), antigen processing and presentation (hsa04612, n = 36), and oxidative phosphorylation (hsa00190, n = 54), where n is the number of TMB-associated genes in each term. Results for all significantly enriched terms are shown in Figure 1—figure supplement 1B, C.

### Principal component analysis

Principal component analysis (PCA) was performed on gene-wise globally z-scored expression matrices using the subset of TMB-associated genes, proteostasis genes, or non-proteostasis genes depending on the analysis. PCA was performed using the sklearn.decomposition.PCA implementation in Python.

### Leave-one-cancer-type-out (LOCO) analyses

To further evaluate whether the inferred TMB-associated transcriptional program was driven by individual cancer types, we performed a LOCO analysis. For each cancer type, gene-level associations with TMB were re-estimated using all remaining cancer types. TMB axis scores for held-out samples were then computed using the same gene set defined from the full-data GLMM, but with the LOCO-derived β. The correlation between TMB axis score and log_10_-transformed TMB was then evaluated within each held-out cancer type.

### Training and held-out validation of the regression coefficient (β)

To assess whether the TMB-associated transcriptional response generalizes to held-out samples, we performed a training/validation split. TMB-associated genes were defined once from the full TCGA cohort and held fixed; only the β values were refit on training data. For each cancer type, 30% of samples were held out as a validation set by stratified random sampling. GLMMs were refit on the remaining 70% using this fixed gene set, and TMB axis scores for validation samples were computed from the held-out β. Correlations between axis scores and log_10_(TMB) were evaluated across all validation samples pooled (pan-cancer).

### Statistical analysis

All statistical analyses were performed using R (v4.5) and Python (v3.9).

## Computational resources

Computing for this project was performed on the Stanford SCG Bioinformatics Cluster (RRID:SCR_026876), owned by the Stanford Genomics Bioinformatics Service Center (RRID:SCR_023340) and operated by Stanford Research Computing (RRID:SCR_023413).

## Acknowledgements

We thank Haiqing Xu and other members of the Petrov lab for their helpful feedback and discussions. This work was supported by the Fellowship from Chan Zuckerberg Initiative to DAP, SCI Innovation Award to DAP and MMW, DOD grant HT9425-25-1-0157 to DAP and NIH grants R01 CA230025 to MMW and DAP, and NIH R35 GM153301 to OB.

## Data Availability

All data used in this study are publicly available.

Tumor RNA-seq gene expression data were obtained from The Cancer Genome Atlas Pan-Cancer (PANCAN) cohort via the UCSC Xena platform, using the batch-corrected dataset EB++AdjustPANCAN_IlluminaHiSeq_RNASeqV2.geneExp.xena (https://xenabrowser.net/datapages/?dataset=EB%2B%2BAdjustPANCAN_IlluminaHiSeq_RNASeqV2.geneExp.xena&host=https%3A%2F%2Fpancanatlas.xenahubs.net), which provides normalized mRNA expression values across TCGA samples with correction for batch effects. Tumor purity estimates and cancer type annotations were obtained from TCGA Pan-Cancer Atlas datasets accessed through UCSC Xena, with purity taken from the ABSOLUTE master calls table (TCGA_mastercalls.abs_tables_JSedit.fixed.txt; https://gdc.cancer.gov/about-data/publications/pancanatlas). Samples were not excluded for missing purity; missing values were imputed as the median within each cancer type.

Tumor mutational burden (TMB) was calculated from the TCGA Multi-Center Mutation Calling in Multiple Cancers (MC3) public MAF (mc3.v0.2.8.PUBLIC.maf.gz; https://gdc.cancer.gov/about-data/publications/pancanatlas; see also https://gdc.cancer.gov/about-data/publications/mc3-2017) by counting missense, nonsense, and silent mutations per sample.

## Code Availability

All code will be made publicly available on GitHub upon publication.

## Notes

### Competing Interest Statement

The authors have declared no competing interest.

