## Supplementary material for "Transcriptional Response to Tumor Mutational Burden Is Consistent Across Cancer Types": Figure 1-figure supplement 1

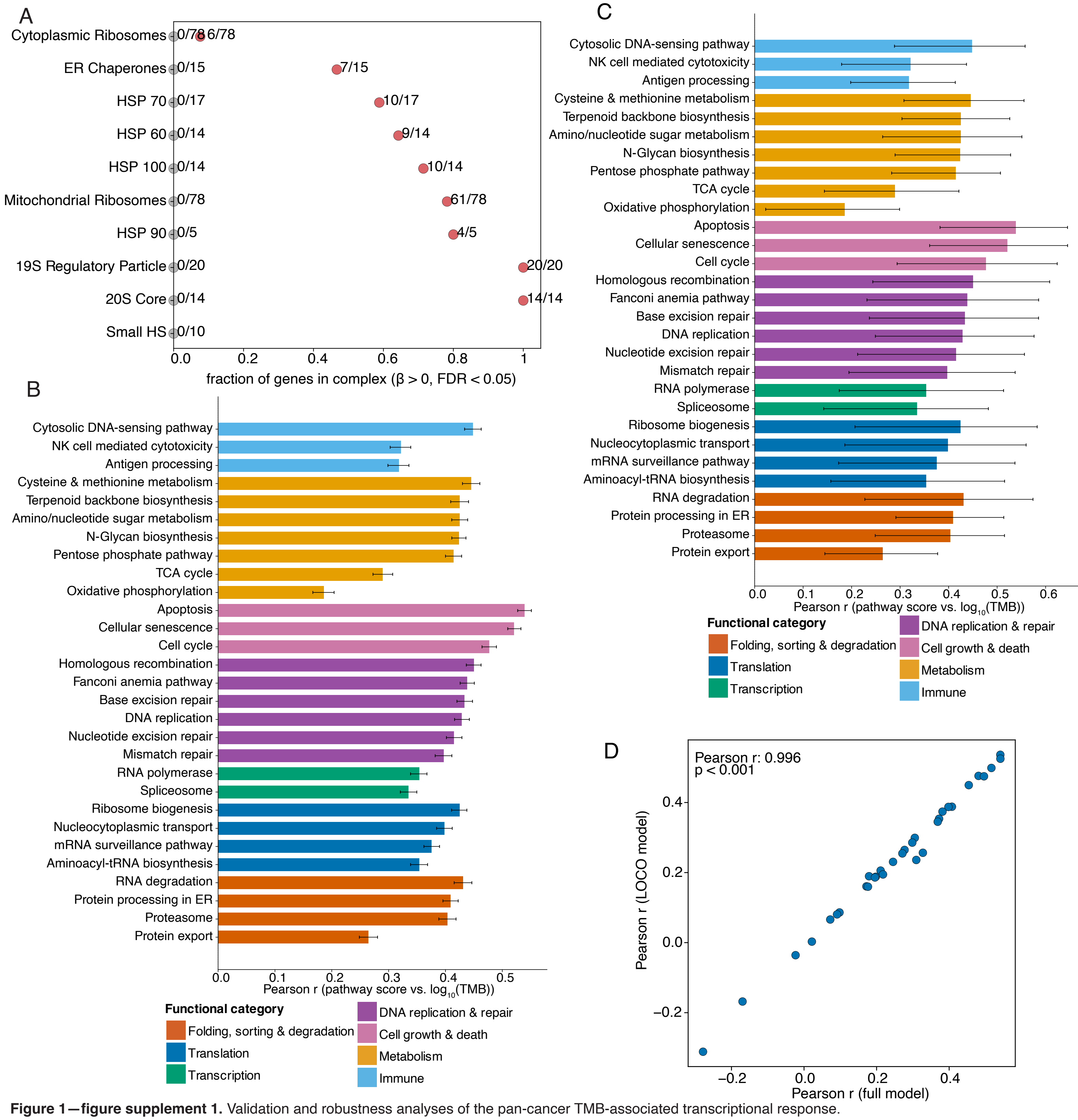


**Figure 1—figure supplement 1.** Validation and robustness analyses of the pan-cancer TMB-associated transcriptional response.
(A) Fraction of genes within each proteostasis complex that are significantly positively associated with TMB (GLMM $\beta > 0$, FDR $< 0.05$). Red points indicate the observed proportion of genes within each complex meeting these criteria. Gray points show the corresponding values from a single permuted dataset, in which $\log_{10}$(TMB) was shuffled within each cancer type. Labels indicate the number of significantly associated genes over the total number of genes in each complex (e.g., 20/20). Proteostasis complexes include ribosomes, proteasome subunits, and chaperones. Proteasome subunits and mitochondrial ribosomes show near-complete enrichment of TMB-associated genes, whereas others, such as the small heat shock (small HS) proteins, show no enrichment, indicating non-uniform engagement of the proteostasis complexes.
(B) Within-cancer-type bootstrap estimates of pathway score–TMB correlations for all significantly enriched KEGG pathways. Pathway scores were computed for each tumor sample and correlated with $\log_{10}$(TMB). All scores were positively correlated with TMB, consistent with the positive TMB associations of the genes used to construct the scores, confirming that the directionality of the transcriptional response is preserved at the pathway level. Error bars represent 95% bootstrap confidence intervals obtained by repeatedly resampling tumors within each cancer type and recalculating the pan-cancer correlation.
(C) Cancer-type bootstrap estimates of pathway score–TMB correlations for all significantly enriched KEGG pathways. The same pathways shown in (B) are shown here. Cancer types were resampled with replacement to assess the stability of pathway score–TMB correlations across different cancer-type compositions. Pathway scores were calculated for each tumor using genes from the indicated pathway. Error bars represent 95% bootstrap confidence intervals from correlations recalculated across all tumors in the resampled dataset for each cancer-type bootstrap replicate.
(D) Robustness of TMB axis score–TMB correlation to leave-one-cancer-type-out analysis. For each cancer type, β coefficients were estimated using all other cancer types (leave-one-cancer-type-out, LOCO). These coefficients were then used to compute TMB axis scores in the held-out cancer type, and the correlation between $\log_{10}$(TMB) and the TMB axis score was calculated. The resulting correlations (y-axis) are compared with those obtained using the full model trained on all cancer types (x-axis). Each point represents a cancer type. The strong agreement (Pearson $r = 0.996$) indicates that the observed associations generalize across cancer types and are not driven by any single dataset.
