## Supplementary material for "Transcriptional Response to Tumor Mutational Burden Is Consistent Across Cancer Types": Figure 2-figure supplement 1

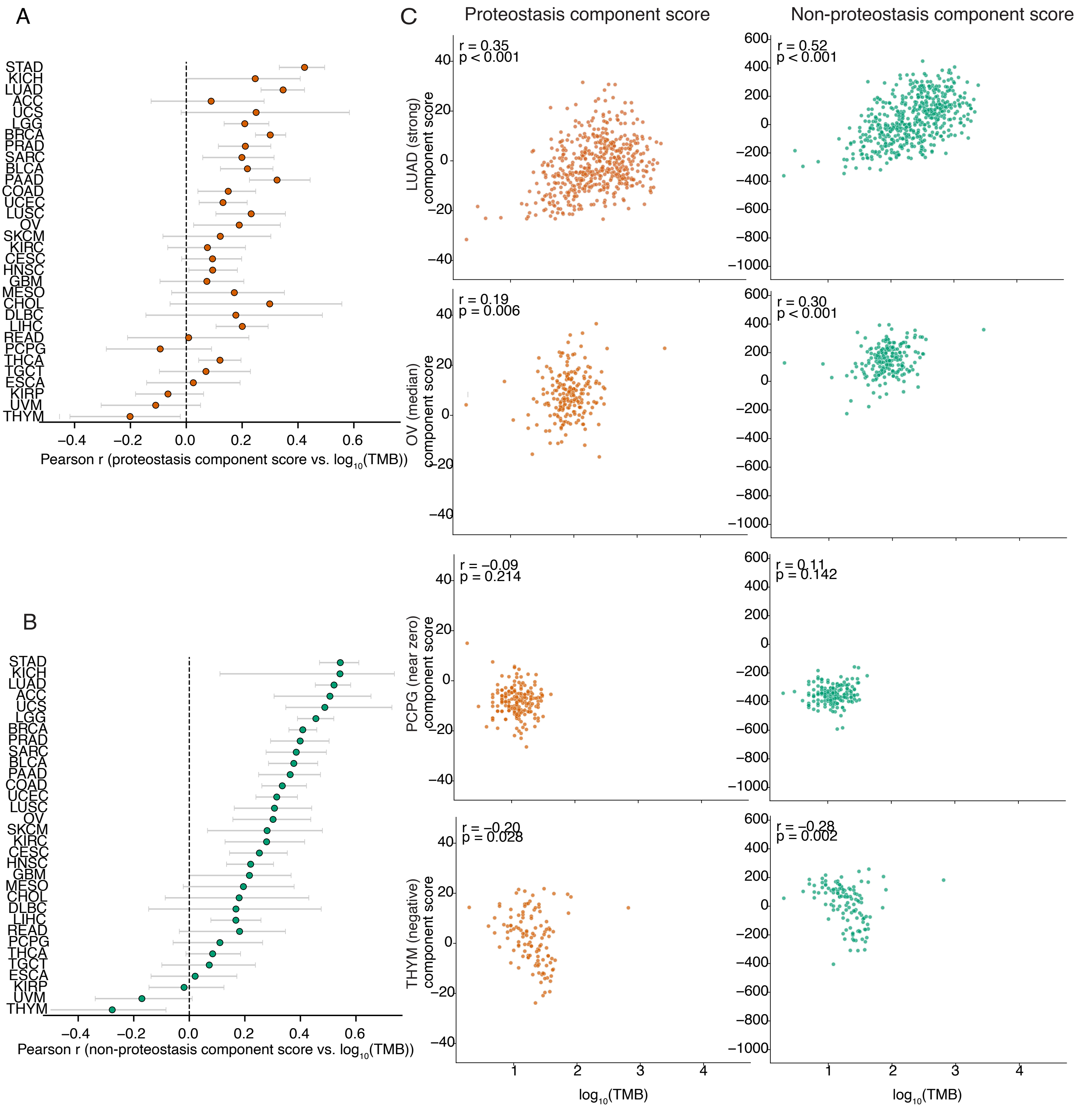


**Figure 2—figure supplement 1.** Transcriptional response to TMB across cancer types in both proteostasis and non-proteostasis components. (A,B) Summary of correlations between $\log_{10}$(TMB) and the proteostasis (A) and non-proteostasis (B) component scores of the TMB axis. Each point represents the correlation within a single cancer type. Error bars represent 95% bootstrap confidence intervals obtained by repeatedly resampling tumors within each cancer type and recalculating the correlation.

(C) Scatter plots of proteostasis (left) and non-proteostasis (right) component scores per tumor. Each row is one cancer type, selected to span a range of TMB axis score–TMB correlation strengths (LUAD strong, OV median, PCPG near-zero, THYM negative). Each dot is a tumor sample. LUAD, lung adenocarcinoma; OV, ovarian serous cystadenocarcinoma; PCPG, pheochromocytoma and paraganglioma; THYM, thymoma.
