## Supplementary material for "Transcriptional Response to Tumor Mutational Burden Is Consistent Across Cancer Types": Figure 3-figure supplement 1

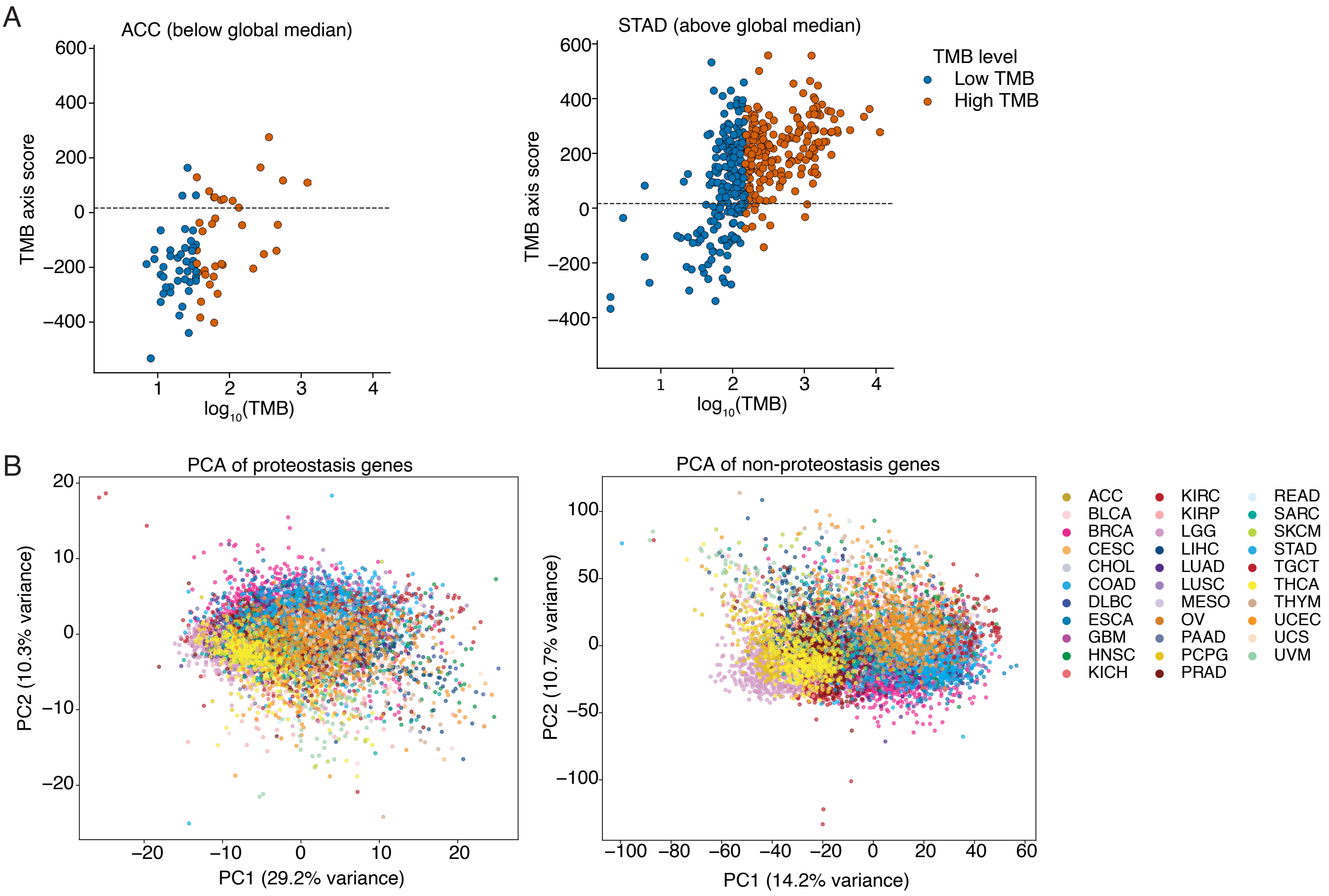


**Figure 3—figure supplement 1.** Baseline gene expression differences across cancer types dominate global transcriptional variation.

(A) Baseline expression varies independently of TMB response across cancer types. ACC and STAD are shown as representative cancer types whose samples cluster below and above the global median TMB axis score, respectively. Despite this difference in baseline expression, both cancer types show a clear positive association between TMB and axis score, illustrating that baseline offsets reflect tissue-specific transcriptional differences rather than differences in TMB response. The dashed line indicates the global median TMB axis score across all TCGA samples. ACC, adrenocortical carcinoma; STAD, stomach adenocarcinoma.

(B) PCA of proteostasis and non-proteostasis TMB-associated genes across TCGA samples. PCA of TMB-associated genes that are (left) proteostasis and (right) non-proteostasis. Colors indicate distinct cancer types. Samples cluster by cancer type in both subsets, indicating that cancer-type-specific transcriptional structure dominates global expression variation regardless of whether genes are proteostasis or non-proteostasis.
